# Codon stabilization coefficient as an index for prediction of codon bias and its potential applications

**DOI:** 10.1101/337634

**Authors:** Rodolofo L. Carneiro, Rodrigo D. Requião, Silvana Rossetto, Tatiana Domitrovic, Fernando L. Palhano

## Abstract

Different methods of mRNA half-life measurements are available, but genome-wide measurements of mRNA half-life in yeast showed a weak correlation between the methods. Moreover, when we compared mRNA half-life determined by these methods with other cellular measurements such as mRNA and protein abundance low correlation was found. To clarify this matter, we analyzed mRNA half-life datasets from nine different groups to determine the most accurate method of measurement. Since codon optimality is one of the significant determinants of mRNA stability, we used the codon stabilization coefficient (CSC) as a reference for mRNA half-life measurement accuracy. After CSC calculation for each dataset, we find strong positive correlations between the CSC from some datasets with other parameters that reflect codon optimality such as tRNA abundance and ribosome residence time. By the use of CSC parameter, we observed that most genes contain non-optimal codons and that codon bias exists toward optimized and non-optimized genes. We also observed that stretches of non-optimal are not randomly distributed since it causes impacts on translation.

## Introduction

mRNA levels are determined by the rates of mRNA synthesis and degradation (Parker, 2012). Genome-wide mRNA half-life measurements showed poor correlation across different datasets and different methods (and, in some cases, even between similar methods), classifying the same mRNA molecule as stable and unstable. It makes the identification of stability sequence motifs and a global view of transcription and translation dynamics in yeast a problematic task(Baudrimont et al., 2017; Harigaya and Parker, 2016). Several features, such as mRNA secondary structure, sequence and structural elements located within 5`and 3`UTR as well as the lengths of the transcripts can affect mRNA stability(Parker, 2012). Herrick and colleagues showed that the percentage of rare codons present in yeast unstable mRNAs were significantly higher than in stable mRNAs(Herrick et al., 1990). Recent experiments on bacteria, yeast, and metazoans indicate that codon optimality is a major determinant of mRNA stability(Boël et al., 2016; Mishima and Tomari, 2016; Presnyak et al., 2015; Radhakrishnan et al., 2016). The concept of codon optimality has been developed through the study of codon bias, which is the frequency at which distinct synonymous codons are present within the genome (Sharp and Li, 1987).

Codon optimality is a term that incorporates the competition between tRNA supply and demand(Pechmann and Frydman, 2012). In several species examined, the cellular tRNA concentrations are proportional to tRNA gene copy numbers(Rocha and Rocha, 2004). Based on this observation, tRNA adaptation index (tAI) calculates codon bias using the genome copy numbers of tRNAs(Dong et al., 1996; Sørensen et al., 1989; Tuller et al., 2010b). The tRNA adaptation index (tAI) reflects the efficiency of the tRNA usage by the ribosome where optimal codons, which are decoded by high abundance tRNA species, are translated more efficiently when compared to non-optimal codons(Reis, 2004). Recently, Coller and colleagues showed a positive correlation between codon optimality and mRNA stability(Presnyak et al., 2015; Radhakrishnan et al., 2016). The authors measured the half-lives of thousands of genes and found that some codons were enriched in the most stable mRNAs while other codons were enriched in the most unstable mRNAs. Based on Pearson’s correlation between the frequency of occurrence of each codon in each mRNA and mRNA half-lives, the authors created the codon stabilization coefficient (CSC) (Presnyak et al., 2015). Direct comparison between codon stabilization coefficient calculated by Coller and colleagues and the tRNA adaptive index (tAI) revealed good agreement between these scores (Presnyak et al., 2015). Nevertheless, as described before, the half-lives of mRNA measured by Coller’s dataset does not correlate well with other mRNA half-lives datasets(Harigaya and Parker, 2016).

We evaluated nine quantitative analyses of mRNA half-lives and calculated the CSC for each dataset. We observed a good correlation of CSC calculated from four studies with different cellular parameters such as tRNA adaptation index, ribosome residence time measured by ribosome profiling and tRNA abundance measured by microarray and RNA seq. Interestingly, these four studies used different methodology to estimate mRNA half-lives. We also measured the CSC average of individual genes (CSCg) using the nine datasets of mRNA half-lives. Again, just four out of nine studies showed good correlation with other genome-wide measurements such as mRNA and protein abundance and frequency of optimal codons. We explored that CSC coefficient to study codon bias, their relation with protein expression and its impact on translation.

## Results and Discussion

### The comparison of global quantification of the yeast mRNA half-lives revels poor correlation among the datasets

In *Saccharomyces cerevisiae*, mRNA half-lives have been globally measured through three main classes of methods, namely, transcriptional inhibition, gene activation control and *in vivo* metabolic labeling (Fig 1A)(Wada and Becskei, 2017). In the transcriptional inhibition method, the RNA polymerase II is inactivated, usually by heat shock (Geisberg et al., 2014; Grigull et al., 2004; Holstege et al., 1998; Presnyak et al., 2015; Wang et al., 2002) and the expression of all genes is reduced. In the gene activation control, the promoter of each gene is substituted (as for GAL or the TET promoter). By controlling the levels of a specific transcriptional activator, it can dissociate from a specific promoter, shutting of the expression of the genes under the control of this promoter(Munchel et al., 2011; Neymotin et al., 2014; Sun et al., 2012). For metabolic *in vivo* labeling, modified nucleotides are introduced into cell media that are incorporated by cells into to the RNA(Baudrimont et al., 2017) the mRNA modified can be recovered by immunoprecipitation or pulled down by streptavidin beads. With nine global quantitative studies of the yeast mRNA half-lives, 5 of which were reported using transcriptional inhibition, three the metabolic *in vivo* labeling and one the gene control method, we observed that the ranges of mRNA half-lives varied substantiated depending on the dataset. For example, in Coller dataset, the mRNA average half-life is 11.5 min (ranging from 0.76 to 60 min), while for Struhl dataset the average is 35.5 min (ranging from 3.4 to 300 min) (Fig 1B). Then we compared the correlation across different datasets and observed a generally poor correlation with Pearson correlation coefficients (r) values ranging from-0.25 to 0.75 (Fig 1C), as reported before(Baudrimont et al., 2017; Harigaya and Parker, 2016). Recently, a comparison among 21 datasets of the yeast proteome measured by mass spectrometry (11 studies), GFP-based studies (9) and TAP-immunoblot (1) showed (r) values ranging from 0.35 to 0.96(Ho et al., 2018). In other words, while for global quantification of the yeast proteome different datasets are convergent, for mRNA half-lives the datasets are divergent (Fig 1C), which hampers the knowledge of a full picture of mRNA turnover in yeast. Relying on any of these datasets is a dubious decision, for we cannot be sure about which one better represents mRNA half-lives. The most accurate mRNA half-life global measurement data should not be necessary following other cellular parameters, such as protein expression, translation efficiency, and mRNA abundance. In fact, when we attempted to identify if one of them meets this criterion, we analyzed all nine mRNA half-life datasets and found little to no correlation between the dataset (mRNA half-life) and the other cellular parameters (protein expression, translation efficiency, and mRNA abundance) (Fig S1). Protein abundance and translation efficiency are controlled by different molecular machinery and may be subjected to additional levels of regulation, what can hinder a direct comparison between mRNA half-life and mRNA or protein abundance. Since we could not find a robust positive correlation on any of the datasets, we needed to find a different method to validate the mRNA half-life measurements.

**Figure 1.**
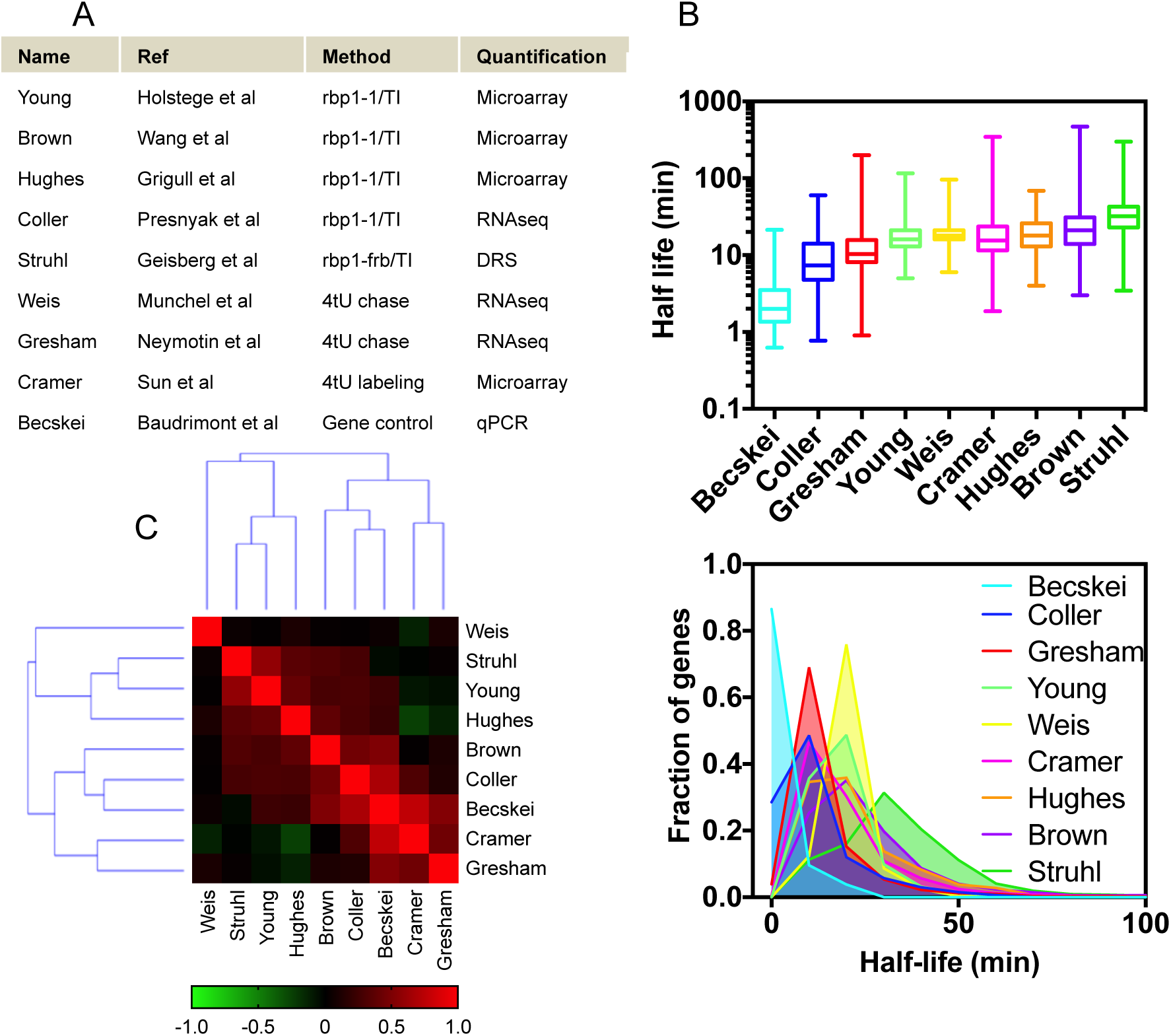
Comparison of different *S. cerevisiae* mRNA half-life datasets. A. Description the measurements and quantification methods from the nine mRNA half-live datasets analyzed namely, Young(Holstege et al., 1998), Brown(Wang et al., 2002), Hughes(Grigull et al., 2004), Coller(Presnyak et al., 2015), Struhl(Geisberg et al., 2014), Weis(Munchel et al., 2011), Gresham(Neymotin et al., 2014), Cramer(Sun et al., 2012) and Becskei(Baudrimont et al., 2017). B. Box plot (upper panel) and frequency of distribution (lower panel) of mRNA half-life measurements in *S. cerevisiae*. C. Spearman correlation between each of the datasets. Values range from-0.25 (negative correlation, green panels) to 1.00 (positive correlation, red panels).

### The codon stabilization coefficient indicates the most biologically accurate mRNA half-life measurements in *S. cerevisiae*

Recent experiments from bacteria(Boël et al., 2016), yeast(Presnyak et al., 2015) and metazoans(Mishima and Tomari, 2016) indicate that codon optimality is a major determinant of mRNA stability. mRNAs can be degraded co-translationally, and codon choice, by influencing the rate of ribosome elongation, affects mRNA half-lives through the action of the DEAD-box protein Dhh1p(Radhakrishnan et al., 2016). The codon stabilization coefficient (CSC) is a correlation between the frequency of occurrence of each codon in mRNA and mRNA half-lives(Presnyak et al., 2015) (Fig 2A). Coller and colleagues created this metric observing the association between optimal codons and mRNA half-lives in their own dataset. We used the nine datasets of mRNA half-lives to create the CSC of each 61 codons in the same way that Coller and colleagues did (Fig. 2A) and then calculated the correlation between these different studies (Fig 2B and S2). We observed that four studies that used different methodologies to quantify mRNA half-lives clustered together, namely, Coller, Becskei, Gresham and Cramer (Fig 2B). This was also evident by a Spearman’s correlation matrix of CSC calculated from all published datasets (Fig 2C). However, it does not mean that these four studies produced the most reliable datasets. The classical metric of optimality of each codon is the tRNA Adaptive Index (tAI)(Pechmann and Frydman, 2012; Sharp and Li, 1987). This metric reflects the efficiency of tRNA usage by the ribosome. When we correlated the CSC score of each codon calculated from published datasets with the tAI of each codon, we observed a similar pattern (Fig 2C). The CSC obtained from Coller, Becskei, Gresham and Cramer’s datasets correlated with tAI with Spearman correlation coefficients (r) ranging from 0.65 to 0.77, while for the other studies the values ranged from-0.69 to 0.23 (Fig 2C). Another metric can be calculated from a powerful methodology used to analyze translation is ribosome footprint profiling (RP). This methodology was created by Jonathan Weissman in 2009 and is based on the deep sequencing of ribosome-protected mRNA fragments (Ingolia et al., 2009). During translation, each ribosome encloses an approximately 30-nt portion of mRNA, protecting it against RNAase digestion. This enclosure allows protected fragments to be sequenced. This approach generates a map, at nucleotide resolution, of translational activity for transcribed genes (Ingolia et al., 2012). We took advantage of the nucleotide resolution of ribosome profiling to see if the difference occupancy at A site for each codon correlates with CSC calculated herein. In principle, if a codon is optimal the ribosome occupancy of this codon must be shorter when compared with a non-optimal codon, so we used 1/ribosome residence time to correlate with CSC (Fig 2C). It is important to note that all ribosome profiling data used herein were obtained in the absence of translational inhibitor, since this treatment distorts ribosome profiles as initiation continue even though elongation is blocked(Gerashchenko and Gladyshev, 2014; Hussmann et al., 2015; Requião et al., 2016; Weinberg et al., 2016). We observed that the CSCs that most agree with three independent experiments measuring codon occupancy time by ribosome profiling(Fang et al., 2018; Gardin et al., 2014; Weinberg et al., 2016) were again the Coller, Becskei, Gresham and Cramer datasets (Spearman correlation coefficients (r) ranging from 0.19 to 0.62). Very recently, it was showed that ribosomal A-site decoding rate impacts normal mRNA decay in yeast reinforcing the correlation among CSC and codon occupancy(Hanson et al., 2018). The pattern observed for codon occupancy time by the ribosome was maintained when other parameters such as tRNA abundance measured by microarray(Tuller et al., 2010) or RNAseq(Weinberg et al., 2016) were used to correlate with CSC (Fig 2C). We also calculated CSC using Spearman correlation instead of Pearson correlation and similar results were found (data not shown). Different from the direct comparison of mRNA half-lives and cellular parameters (Figure S1) comparison with the datasets CSCs gives us strong positive correlations, meaning that we are able to identify the datasets that are most coherent with other parameters that are directly involved in codon adaptability and therefore have an immediate effect in mRNA half-life. These data together suggest that four datasets, Coller, Becskei, Gresham and Cramer, are the most reliable, since their CSCs agree with different experimental data, while the others do not, moreover, the correlation among these studies showed (r) ranging from 0.12 to 0.75 (Fig 1C) and the correlation among the CSC calculated from these studies ranged from 0.78 to 0.91 (Fig 2C).

**Figure 2.**
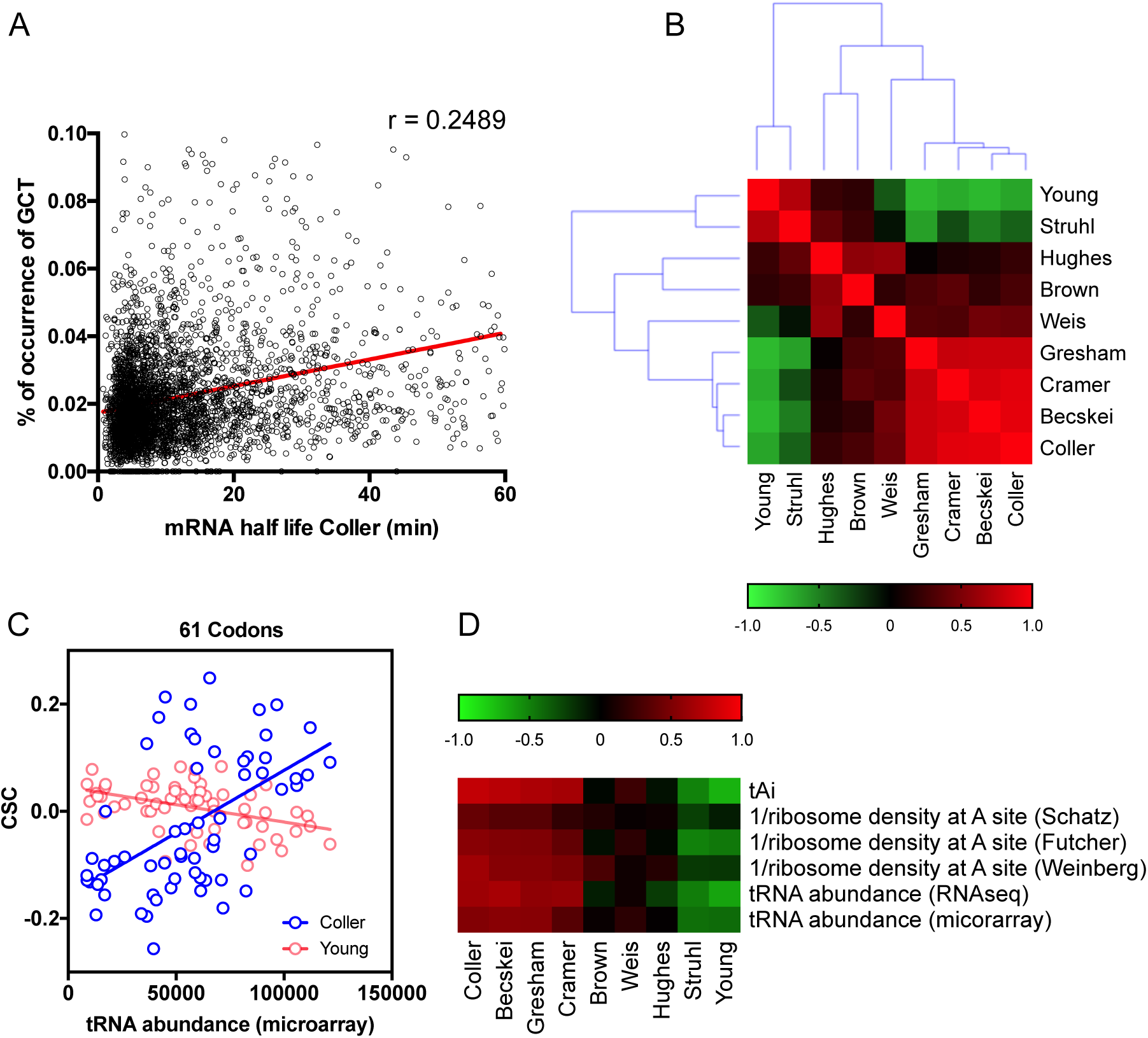
The codon stabilization coefficient (CSC) as a metric of mRNA half-life measurement accuracy. A. Codon stabilization coefficient of the codon GCT based on Collers’s group mRNA half-life measurement. The Person correlation coefficient r determines CSC. B. Pearson’s correlation between CSCs of each of the datasets for the 61 yeast codons. Values range from-0.64 (negative correlation, green panels) to 1.00 (positive correlation, red panels). C. Correlation between CSCs for the 61 yeast codons calculated using Coller’s and Young’s mRNA half-life with tRNA abundance (Tuller et al., 2010a). Spearman’s correlation was 0.51 and-0.42 for Coller’s and Young’s CSC respectively. D. Spearman’s correlation between CSCs for the 61 yeast codons from each of the datasets with different cellular parameters; tAI(Sharp and Li, 1987), 1/ribosome density at A site(Fang et al., 2018; Gardin et al., 2014; Weinberg et al., 2016) and tRNA abundance(Tuller et al., 2010; Weinberg et al., 2016). Values range from-0.72 (negative correlation, green panels) to 0.7 (positive correlation, red panels).

However, if one is interested to know the yeast genes mRNA half-life absolute values, Coller, Becskei, Gresham and Cramer studies can give entirely different answers. In other words, a good correlation between two studies can hide essential differences in the range of mRNA half-life absolute values obtained by each research group. This is probably an inevitable consequence of using different experimental methodologies.

### Mean values of CSC computed for each gene

Another way to estimate the biological significance of the CSC is the analysis of the correlation of the mean values of CSC computed for each gene with other genome-wide measurements apart from mRNA half-lives. When we measured the CSC average of individual genes (CSCg) using Coller’s dataset, we observed an almost perfect correlation with the frequency of optimal codons calculated by Saccharomyces genome database (SGD) (Fig 3A). Agreement of CSC with SGD database further strengthens the CSCs as an indicator of mRNA half-life measurements accuracy. Next, we did the same analysis using the CSCs calculated from the other eight mRNA half-lives datasets and compared with the frequency of optimal codons, protein abundance(Lahtvee et al., 2017), mRNA abundance(Yassour et al., 2009) and transcriptional elongation(Weinberg et al., 2016) (Fig 3B). We noticed that the CSCg calculated from Coller’s dataset was better correlated with all biological parameters analyzed when compared to other CSCs obtained from other datasets (Fig 3B). In that way, in the subsequent experiments, we chose to continue this study only with Coller’s CSC.

**Figure 3.**
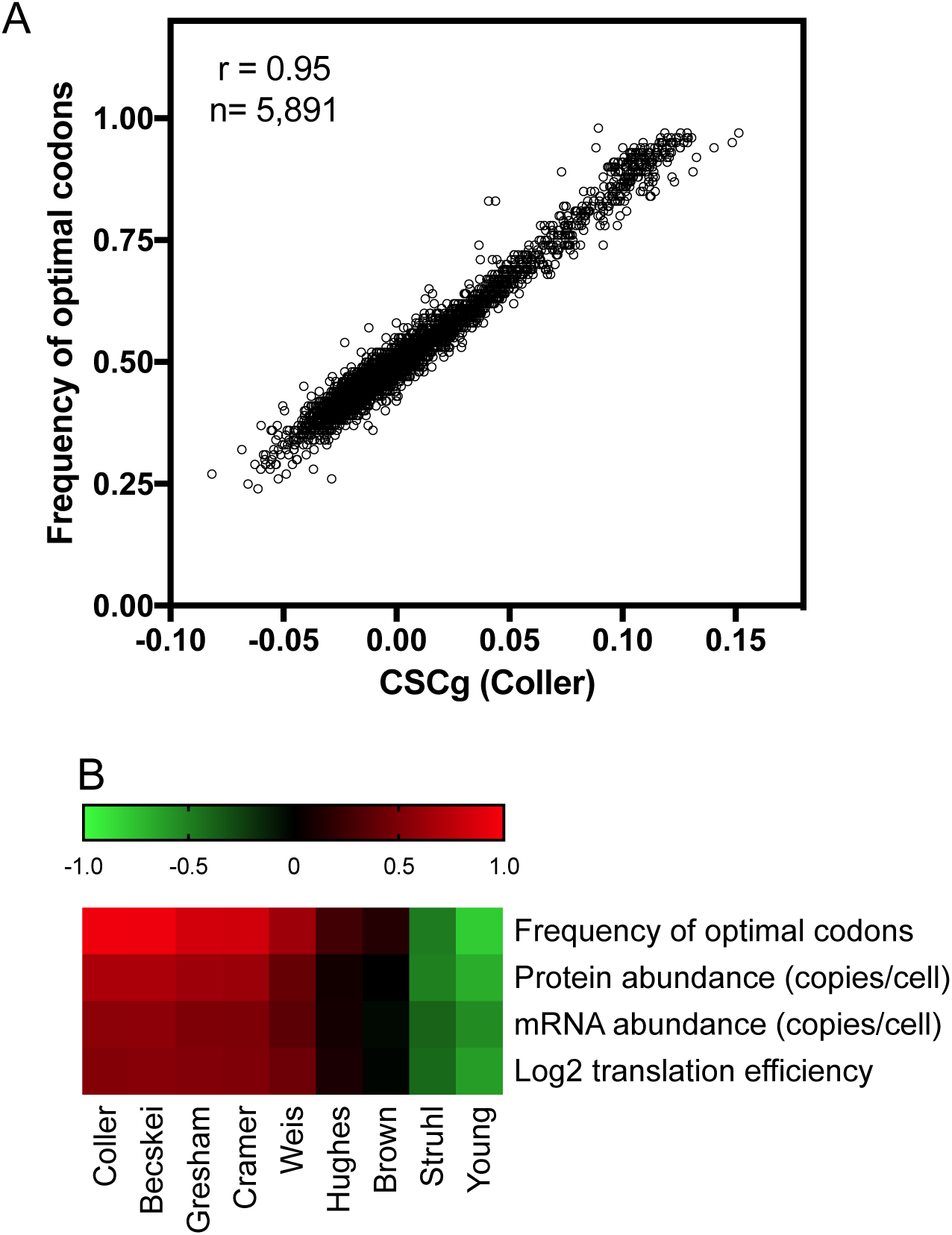
Individual genes CSC (CSCg) correlation with different cellular parameters. The CSC for each of 61 yeast codons was calculated using the nine mRNA half-life dataset (Figure 2B). The CSC calculated for every 61 codons were used to calculate a mean for each of 5,891 yeast genes generating the CSCg. A. A positive correlation between Coller’s dataset CSCg and Saccharomyces genome database frequency of optimal codons (r = 0.95). B. Spearman’s correlation between CSCg calculated from all nine mRNA half-lives datasets with the frequency of optimal codons (Saccharomyces Genome Database), protein abundance(Ho et al., 2018), mRNA abundance(Yassour et al., 2009) and translation efficiency(Weinberg et al., 2016). Values range from-0.79 (negative correlation, green panels) to 0.95 (positive correlation, red panels).

Next, we analyzed the distribution of all yeast genes taking in to account their CSCg. We can observe that the majority of genes present a non-optimal CSCg (Fig 4A, black circles). A more precise analysis revealed that two thirds of the CSCgs (66%) are non-optimal (Fig 4A, inset, red line), while one third (34%) is optimal (Fig 4A, inset, blue line), with approximately 4% of the genes presenting a high level of optimality (Fig 4A, inset, dark blue line). To determine if the codons bias acts just in the small fraction of genes enriched in optimal-codons, we created a scramble yeast genome. We scrambled all codons, but we maintained the protein sequence of all ORFs, as well as the proportion of all 64 codons from the full genome. The CSCg distributions of scrambled genes showed that random codon choice creates a uniform genome where most genes possess neutral CSCg (Fig 4A, blue squares). Interestingly, we also observed a codon bias toward genes with extremely low CSCg. One possibility to explain this observation is that a small fraction of genes concentrates the majority of optimal codons with the highest CSC scores and, consequently, most genes use codons with lower CSC scores, increasing the chance to have genes with a high amount of non-optimal codons. To address this point, we analyzed just the genes with CSCg lower than 0 (3,258 genes), and we scrambled the codons of these genes. As observed in Figure 4B, the frequency of genes with extremely low CSCg was ten times higher in the yeast genome when compared with the scrambled genome. We concluded that codon bias exists as a force to select genes with a high concentration of both optimal and non-optimal codons.

**Figure 4.**
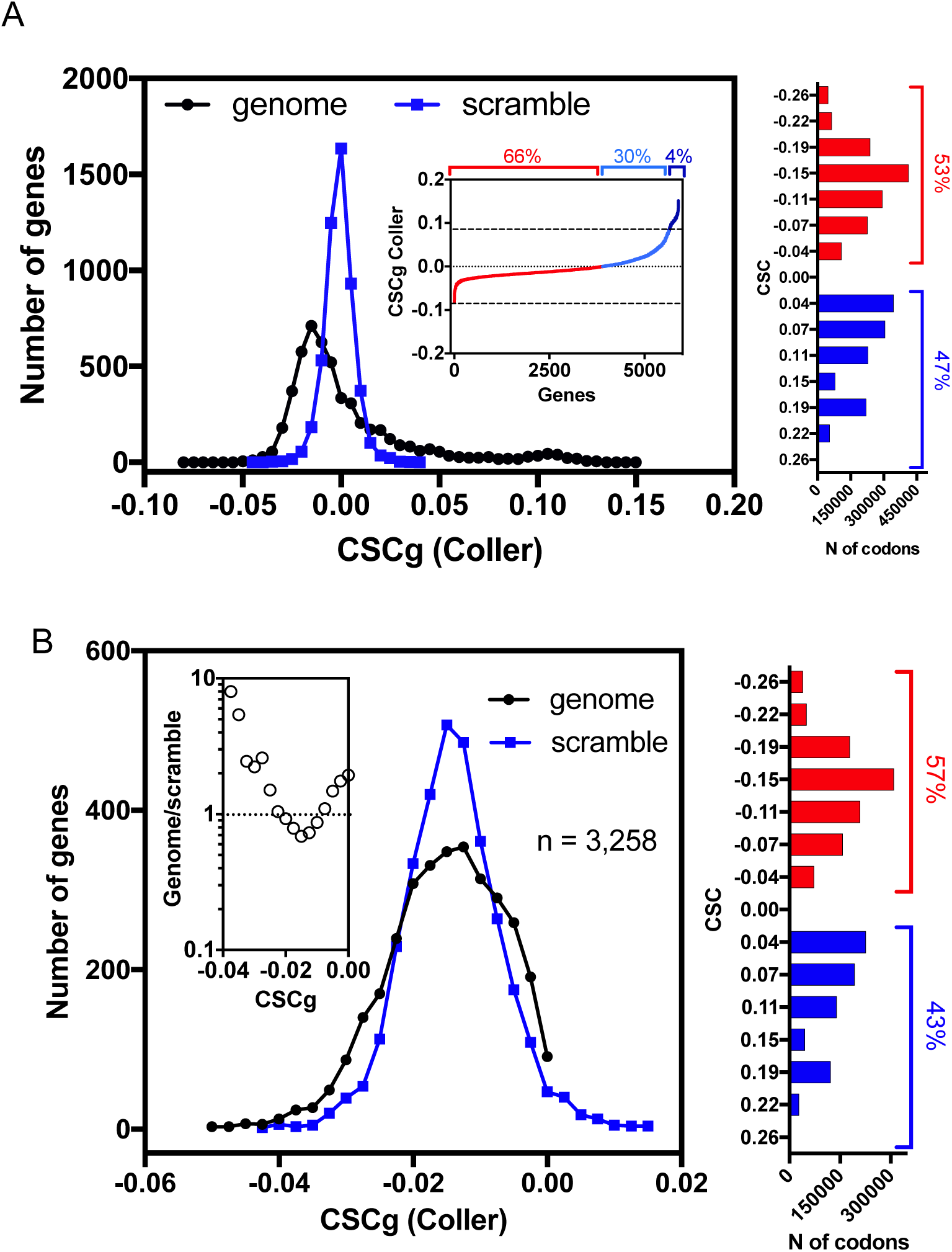
Codon bias acts on both optimal and non-optimal codon concentration. A. Distribution of *S. cerevisiae*’s genome (black line) and scrambled genome (blue line) CSCg (Coller). In genome, 66% of the genes possess negative CSCg, while 34% possess positive CSCg (inset). B. Distribution of *S. cerevisiae*’s genome (black line) and scrambled genome (blue line) genes with CSCg lower than 0. The frequency of genes with CSCg lower than 0 is ten times higher in the *S. cerevisiae*’s genome when compared with the scrambled genome (inset). The frequency of distribution of codons based on CSC score of the full yeast genome (panel A) or the just the genes with CSCg equal or lower than 1 (panel B) is showed in the left.

### Highly expressed genes concentrate the optimal codons in yeast

When we correlated the CSCg with the reads per kilobase per million (measured for each gene by ribosome profiling(Heyer and Moore, 2016)), we observed a positive correlation (r = 0.63) (Fig 5A). The 61 codons were then arbitrarily classified into three categories based on CSC. We considered codons with CSC higher than 0.1 optimal-codons, while codons with CSC lower than-0.1 were non-optimal codons (Fig 5B). The codons with CSC in the range of-0.1 to 0.1 were considered neutral codons (Fig 5B). Using the criteria described above, we divided all yeast genes arbitrarily into ten groups (Fig 5C). After k-means cluster we observed three main categories (Fig 5C). The first category (cluster 1) was comprised of genes composed by codons with optimal or neutral scores and virtually none of the non-optimal codons. Even though less than 6% of the genes were found in this category, they were responsible for approximately 70% of all transcripts in the cell (Fig 5C). Genes characterized in the second category (cluster 2) have a high abundance of neutral codons and represents 26% and 20% of the number of genes and transcripts in the cell respectively (Fig 5C). Finally, the third category (cluster 3) was the inverse of the first, and virtually none of the optimal codons were found. Most genes (62%) fall in this category, but they represent just 10% of all transcripts in the cell (Fig 5C). Even though a small number of genes are responsible for most of the transcripts, they are enriched with optimal codons, confirming the relationship between codon optimality and higher levels of translation.

**Figure 5.**
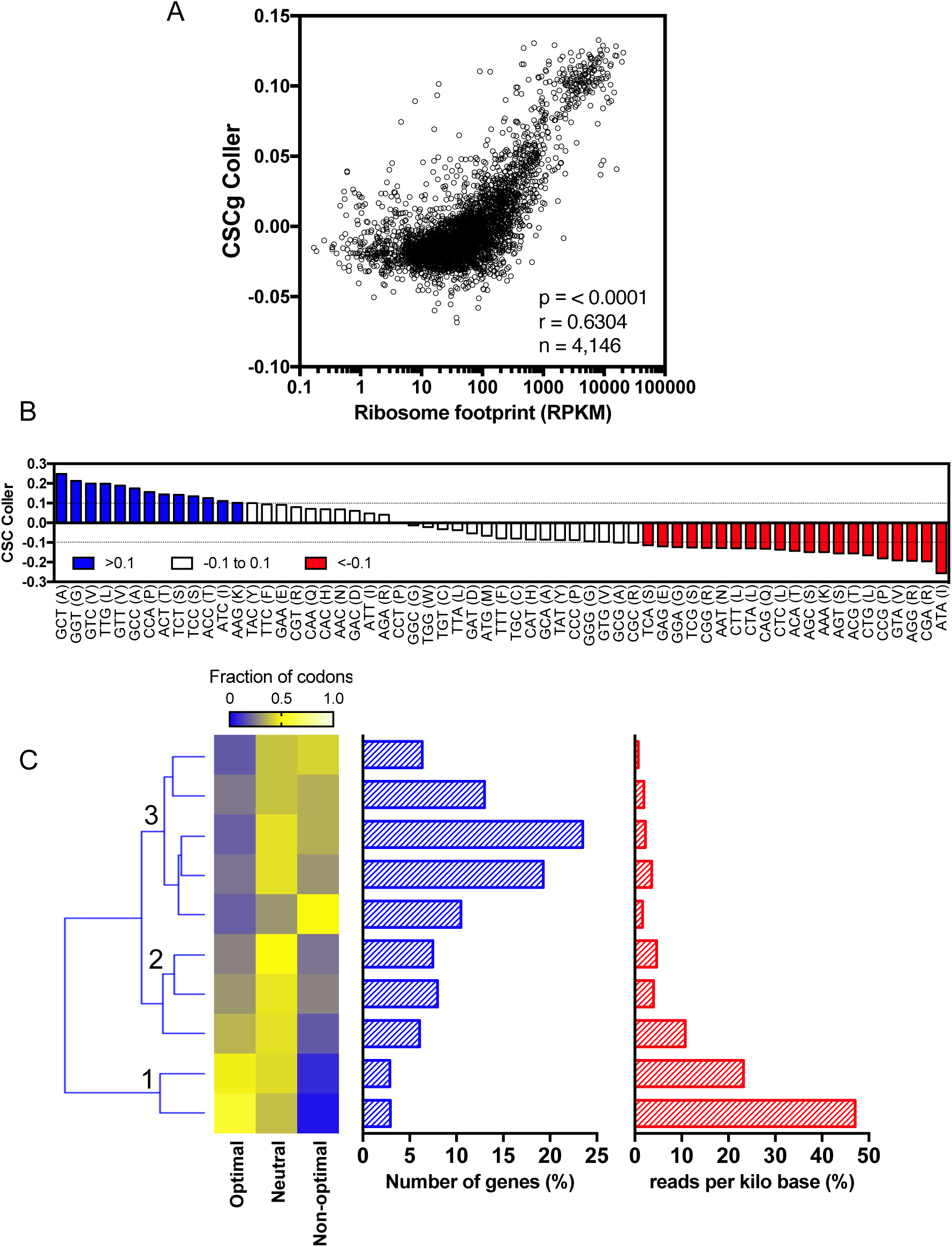
Correlation between CSC and the number of transcripts in *S. cerevisiae*. A. Positive correlation between CSCg and the reads per kilobase per million (r = 0.63) (Heyer and Moore, 2016). B. CSC plotted for each codon with optimality represented by different colors. Optimal codons (CSC > 0.1) in blue, neutral codons (0.1 > CSC >-0.1) in white and non-optimal codons (CSC>-0.1) in red. C. k-means cluster of *S. cerevisiae* genes based on optimal, neutral and non-optimal codons. The number of genes in each cluster is represented by the blue bars, while the number of transcripts is represented by the red bars.

### Correlation between CSCg and the copy number of proteins in yeast

As described in Figure 3B, we observed a positive correlation between CSCg from Coller’s dataset and protein abundance. We analyzed this correlation with more details in Figure 6A(Breker et al., 2013). Besides the Pearson correlation coefficients (r) of 0.66, we noticed the presence of outliers in the left superior quarter of the plot (blue points, Fig 6A). All proteins with more than 10^5^ copies / cell have CSCg higher than 0. Nevertheless, some proteins with the highest CSCg (>0.1) are low abundant (less than 10^3^ copies/cell). To explain this discrepancy, we compared the fold of induction / repression in more than 400 conditions of inliers *vs* outliers (Fig 6B). It is clear that outliers are genes more inducible when compared to inliers. It means that these genes possess high CSCg because in some moments it is essential for the cells to produce high amounts of mRNA and proteins to deal with a new environmental or physiological condition. To support this hypothesis, we calculated the CSCg of genes up-regulated in different stress conditions (Fig 6C and 6D). We used as threshold 1.5 fold induction, as represented in Figure 6C when the cells were exposed to DTT(Breker et al., 2013). We observed that, for all stress conditions analyzed, the CSCg of induced genes were higher when compared with of the non-induced genes (Fig 6C), confirming our hypothesis that high CSCg values are related to stable mRNA and abundant protein expression, under certain conditions.

**Figure 6.**
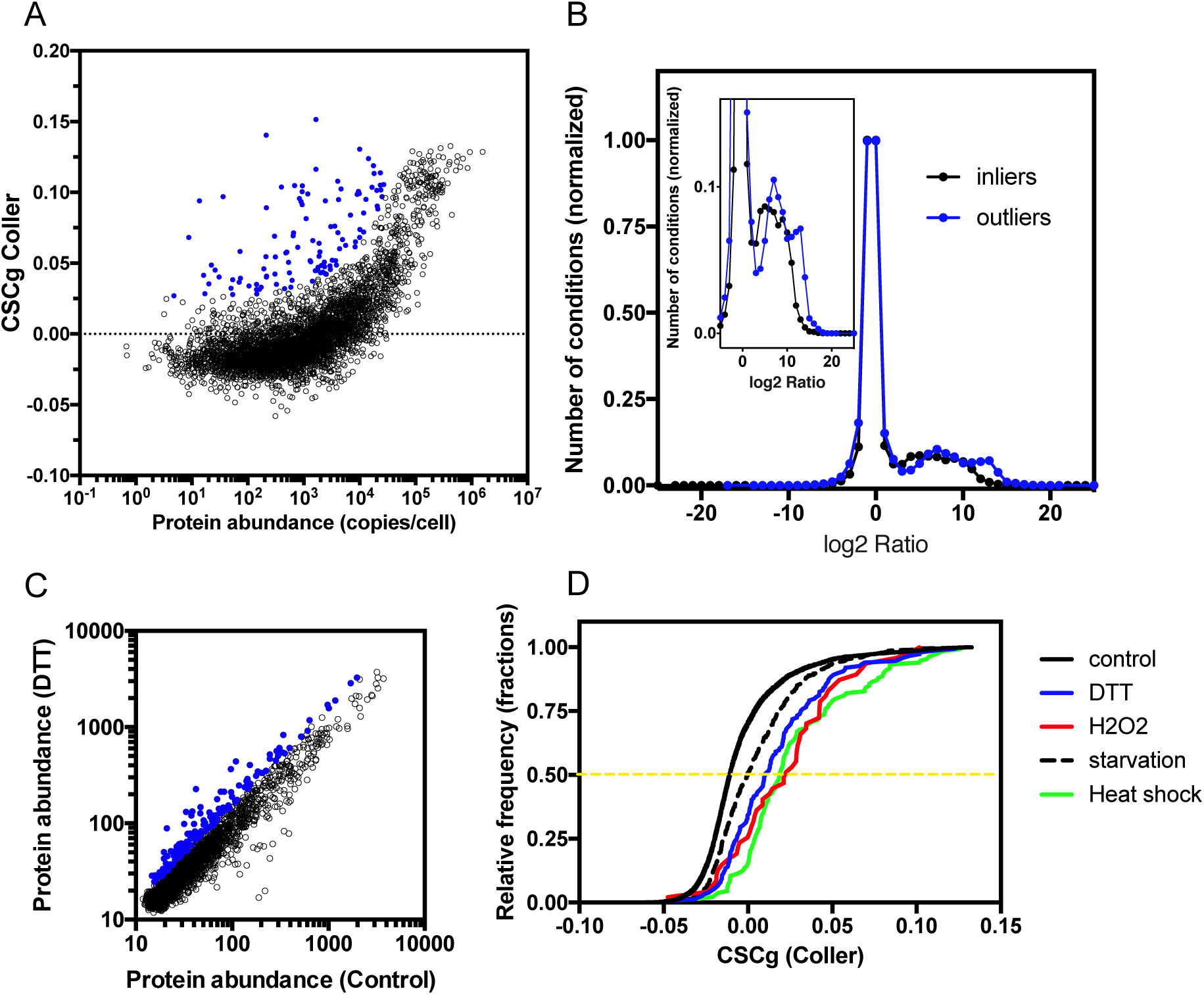
Correlation between CSCg and the copy number of proteins in *S. cerevisiae*. A. Positive correlation between CSCg and protein abundance (r = 0.66) (Kulak et al., 2014). Outliers are highlighted in blue. B. The level of induction/repression at 400 different experimental conditions (data obtained from SGD) was compared using the inliers genes (black line) and outliers (blue line) genes of panel A. The inset is the magnification of the panel B. It is clear that the outliers genes of panel A achieve, more often, higher levels of induction when compared to control (inliers). C. Induced proteins (blue dots) expressed 1.5 fold higher after cellular exposure to DTT when compared to non-stressed cells(Breker et al., 2013). D. The CSCg of genes induced 1.5 fold at different stresses (Breker et al., 2013) were compared to all genes (control). For all stresses analyzed the CSCg of genes overexpressed are higher when compared to the control.

### Distribution of codons in the yeast genome and its impact in translation

Previous reports showed that non-optimal codons are concentrated in both 3’and 5’regions in several genes(Tuller et al., 2010; 2011). The biological explanation for this observation relays in the efficiency that ribosome translates these codons(Tuller et al., 2010). The ribosome ramp hypothesis claims that highly expressed genes possess, in their 3’mRNA regions, a subset of non-optimal codons, which would be necessary to slow down translation, thus avoiding ribosome jams and minimizing the cost of protein translation. However, the ribosome profiling data used to support the ribosome ramp hypothesis was conducted in the presence of cycloheximide before cell lysis. As mentioned before, this protocol disrupts the data, especially in the region nearby the ATG(Gerashchenko and Gladyshev, 2014; Hussmann et al., 2015; Weinberg et al., 2016). Using the CSC as a metric, we measured the occupancy of non-optimal codons in the yeast genome and, as observed before by the use of tAI (Tuller et al., 2010), we observed an enrichment of non-optimal codons from codon 3 until codon 10 (Figure 7A). A more discrete enrichment was also observed in the region close to the stop codon (Figure 7A).

**Figure 7.**
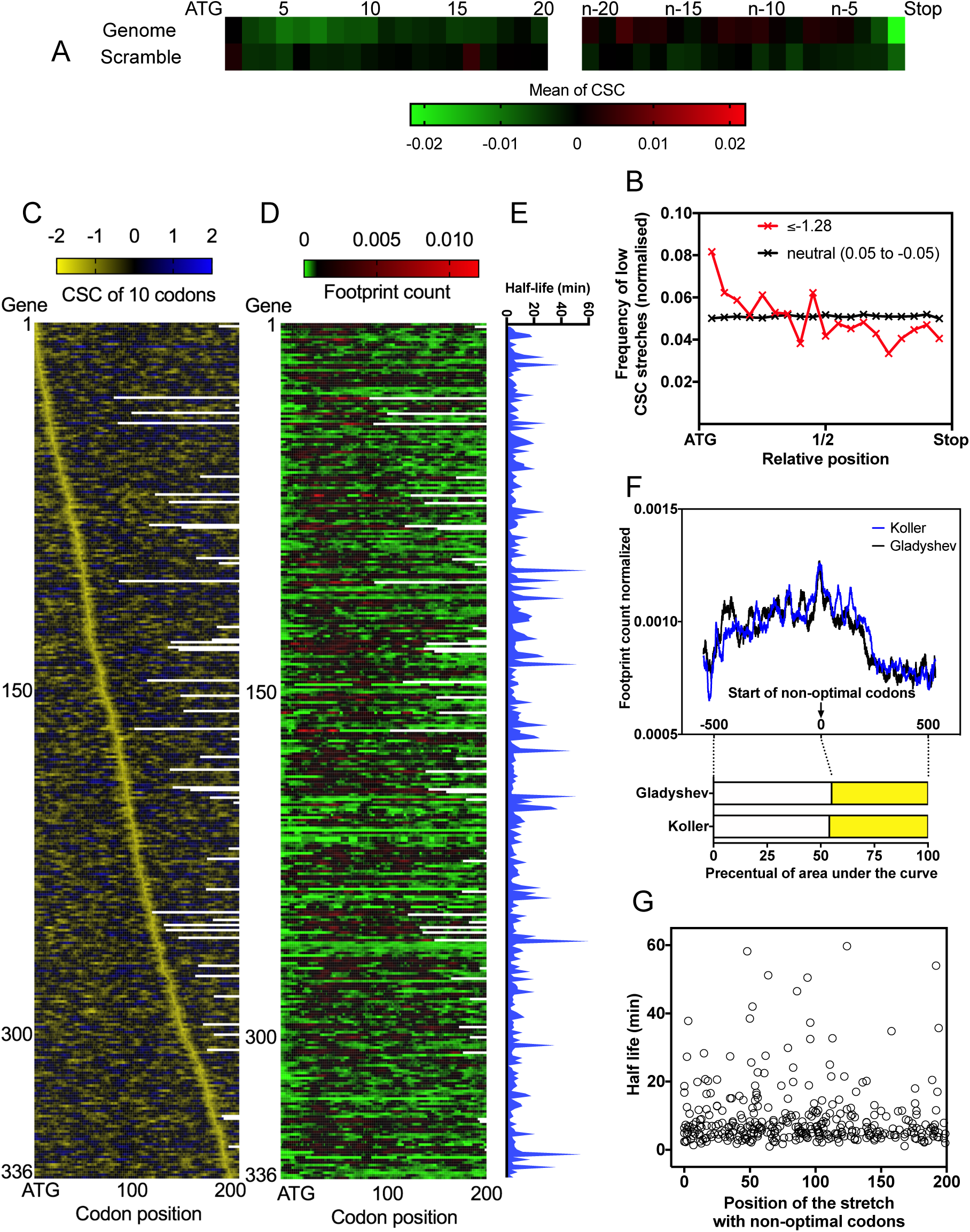
Distribution of codons in yeast genome and its impact in translation. A. Enrichment of low CSC codons near both the N-and C-terminals. Scrambled genome does not present low CSC codon enrichment. The numbers 0.02 and-0.02 are the mean of CSC for each codon from the full yeast genome. The left part of the panel B represents the first 20 codons while in the right are represented the last 20 codons. B. *S. cerevisiae*’s genome displays an enrichment of stretches of ten codons with extremely low CSC (≤-1,28) (red line) near the ATG. Neutral stretches (0.05 ≥ CSC ≥-0.05, black line) are distributed equally in gene and was used as a control group. C. Alignment of 336 *S. cerevisiae* genes (first two hundred codons) based on the position of their low CSC stretches of ten codons (≤-1,28) in relation to the ATG, from closest (top) to furthest (bottom). The numbers-2 and 2 are the sum of CSC in a window of ten codons. Normalized footprint reads (D) and mRNA half-lives (E) of the genes presented at panel 7C. The numbers 0.005 and 0.010 represent the normalization of ribosome profiling analyzes. The normalization was performed by dividing the number of reads at each position by the total number of reads in each gene. F. Average ribosome footprint density of the genes presented in panel 7D aligned based on the stretch of non-optimal codons (position 0). The area under the curve of the 500 nucleotides before the non-optimal stretch of codons (−500 to 0, white bar) was compared with the area under the 500 nucleotides after the stretch (0 to 500, yellow bars). Two independent ribosome profiling studies were used (Gerashchenko and Gladyshev, 2014; Pop et al., 2014). G. No correlation of mRNA half-life measurement (Presnyak et al., 2015) with the position of the stretch with non-optimal codons (CSC ≤-1,28) was observed.

Recently, it was showed that the DEAD-box protein Dhh1p was a sensor of codon optimality that targets mRNA for degradation(Radhakrishnan et al., 2016). In a series of elegant experiments, Coller and colleagues showed that Dhh1p binds to ribosomes and modulates ribosome occupancy of mRNAs with low codon optimality(Radhakrishnan et al., 2016). The authors proposed that Dhh1p binds to translating ribosomes when elongation is slow and that the number of slow-moving ribosomes stimulates mRNA decay. To prove that, a gene with optimal codons (PGK1) was modified with a non-endogenous stretch of ten codons with exceptionally low CSC (TTA ATA GCG CGG CGG CGG CGG CGG GCG ACG). This stretch was placed at increasing distance from the ATG. Interestingly, the half-live of PGK1 mRNAs correlated inversely with the distance of which the stretch was placed, in other words, mRNAs with non-optimal codons concentrated close to ATG possess higher half-lives when compared to RNAs where these stretches are further apart(Radhakrishnan et al., 2016). We asked if this observation could be expanded to endogenous mRNAs. Using the CSC score we sum the value of each one of ten codons used in Coller’s constructs and we found a value of - 1.28. To determine the frequency and the position of stretches of non-optimal codons in yeast genome, we developed a program that is able to screen the sequence of a given gene and calculate the sum of CSC of every 10 codons. We found 1,700 stretches present in 1,200 genes with values equal or lower than-1.28. The distribution of these stretches was not random, they were concentrated close to the ATG (Figure 7B). We used the 1,200 genes with values equal or lower than - 1.28 in a window of ten codons and removed the genes with more than one stretch. After that, we took just the genes where the half-life of mRNA was calculated by Coller’s lab. The remaining genes where aligned in a window of 200 codons in a crescent order of position of a stretch with non-optimal codons (Figure 7C). Next, the ribosome profiling of the same genes was analyzed (Figure 7D). It is important to note that we used two independent ribosome profiling data (Koller and Gladyshev) both performed in the absence of cycloheximide before cell lysis(Gerashchenko and Gladyshev, 2014; Pop et al., 2014). We clearly observed increased ribosome density associated with the stretch of non-optimal codons (Figure 7D and 7F). We expected to see a high density of ribosomes before the stretch with non-optimal codons (Figure 7F). In fact, the ribosome density present in a window of 500 nucleotides before the stretch of non-optimal codons was higher when compared with the same window of nucleotides after the stretch (Figure 7D and 7F). Based on Coller’s data the genes with non-optimal stretch of codons closed to ATG should have higher half-life when compared to genes where the stretch is further apart, but it was not the case (Figure 7E), no correlation was observed (Figure 7G). In other words, the presence of non-optimal codons was sufficient to induce ribosome stalling (Figure 7D) and to slow ribosome movement before the non-optimal codons had been reached (Figure 7F) but their position in mRNA didn’t influence the half-life of mRNA (Figure 7E and 7G).

### Conclusions

We conclude that the CSC is a useful parameter for estimate mRNA half-lives based on codon profile. We suggest the use of CSC as a parameter to calculate the accuracy of wide measurement of mRNA decay. Moreover, using CSC of each 61 codons as well the CSCg calculated for each yeast gene we were able to understand important features of yeast genome such as; 1) most genes contain non-optimal codons, 2) the codon bias exists toward optimized and non-optimized genes, 3) genes with optimal codons usually are more translated and the exceptions are translated in stresses conditions, 4) stretches of non-optimal codons are concentrated closed to the ATG in yeast transcripts but an explanation for this observation still missing.

## Methods

| Reagent or resource | Source | Identifier |
| --- | --- | --- |
| tAI | Sharp and Li, 1987 |  |
| Ribosome density at A site | Fang et al., 2018; |  |
| Ribosome density at A site | Gardin et al., 2014 |  |
| Ribosome density at A site | Weinberg et al., 2016 |  |
| tRNA abundance | Tuller et al., 2010 |  |
| The frequency of optimal codons | Saccharomyces Genome Database |  |
| Protein abundance | Ho et al., 2018 |  |
| mRNA abundance | Yassour et al., 2009 |  |
| Reads per kilobase per million | Heyer and Moore, 2016 |  |
| Protein abundance | Kulak et al., 2014 |  |
| Induced stress proteins | Breker et al., 2013 |  |
| *S. cerevisiae* CHX-free Riboseq data | Gerashchenko and Gladyshev, 2014 | SRR1520311 |
| *S. cerevisiae* CHX-free Riboseq data | Weinberg et al., 2016 | SRR1049521 |
| *S. cerevisiae* CHX-free Riboseq data | Pop et al., 2014 | SRR1688545 |
| **Software and algorithms** |  |  |
| Codon stabilization coefficient gene (CSCg) | This paper |  |
| Scrabble genome | This paper |  |
| Localization of non-optimal codons stretches | This paper |  |

### Data sources

Coding sequences and annotation of *S. cerevisiae* were obtained from Saccharomyces cerevisiae genome database and Ensemble Genomes. We excluded 746 dubious ORFs and mitochondrial all ORFs, as defined in the Saccharomyces Genome Database, from our analysis. We gathered 9 datasets from published studies measuring mRNA half-life in yeast, Young(Holstege et al., 1998), Brown(Wang et al., 2002), Hughes(Grigull et al., 2004), Coller(Presnyak et al., 2015), Struhl(Geisberg et al., 2014), Weis(Munchel et al., 2011), Gresham(Neymotin et al., 2014), Cramer(Sun et al., 2012) and Becskei(Baudrimont et al., 2017).

### Codon stabilization coefficient (CSC) and codon stabilization coefficient gene (CSCg) calculation

We calculated the CSC of each 61 codons as described previously (REF). To calculate the codon stabilization coefficient in a given gene (CSCg), a program that can screen the coding sequence of a given gene and attribute different values to different codons was developed. The program sums the CSC values of all codons of a gene and divides by the number of codons present in this gene.

### Ribosome profiling data

To analyze the correlations between the ribosome density and the net charges, *S. cerevisiae* ribosome profiling data were used (REF). The data were analyzed as described previously (REF). Briefly, the data were downloaded from GEO, and the adaptors (CTGTAGGCACCATCAAT) were trimmed. The trimmed FASTA sequences were aligned to *S. cerevisiae* ribosomal and noncoding RNA sequences to remove the rRNA reads. The unaligned reads were aligned to the *S. cerevisiae* S288C genome, which is deposited in the Saccharomyces genome database. First, any reads that mapped to multiple locations were removed. Then, the reads were aligned to the *S. cerevisiae* coding sequence database, allowing two mismatches per read. Genes with <50% of positions covered were eliminated.

### Clustering Analysis

Clustering analyses were performed by Euclidean distance using the software Orange3. Clustering by k-means was performed with ten defined centers using the software Orange3.

### Scrabble genome

A program that maintains the ORF length as well the amino-acid sequence generated by the scrabble genome. The program scrabbled the codons at a given genome keeping the proportionality of the codons of the input genome.

### Localization of non-optimal codons stretches

We created a program that calculates the sum of CSC of each codon in a determined window for each ORF. We defined a value of CSC of-1.28 in a window of ten codons that reflects the sum of CSC of the codons TTA ATA GCG CGG CGG CGG CGG CGG GCG ACG. The insertion of these codons in a gene causes a destabilization on mRNA half-life (REF). The program sums the CSC of ten codons and when values equal o lower than-1.28 was found the program localized the position of the non-optimal codons in the ORFs.

## Acknowledgments

We thank Maite Vaslim, Edson Delatorre, and João Trindade Marques for critically reading the manuscript.

## Author contribution

Conceptualization, F.L.P.; Methodology, R.L.C., R.D.R., S.R., T.D., and F.L.P.; Investigation, R.L.C., R.D.R., T.D., and F.L.P.; Writing, F.L.P.; Funding Acquisition, S.R., T.D., and F.L.P.; Resources, T.D., and F.L.P.; Supervision, T.D., and F.L.P.

